# Sex estimation in cranial remains: A comparison of machine learning and discriminant analysis in Italian populations

**DOI:** 10.1101/2020.04.30.071597

**Authors:** A Pozzi, C Raffone, MG Belcastro, TL Camilleri-Carter

## Abstract

**Objectives:** Using cranial measurements in two Italian populations, we compare machine learning methods to the more traditional method of linear discriminant analysis in estimating sex. We use crania in sex estimation because it is useful especially when remains are fragmented or displaced, and the cranium may be the only remains found.

**Materials and Methods:** Using the machine learning methods of decision tree learning, support-vector machines, k-nearest neighbor algorithm, and ensemble methods we estimate the sex of two populations: Samples from Bologna and samples from the island of Sardinia. We used two datasets, one containing 17 cranial measurements, and one measuring the foramen magnum.

**Results and Discussion:** Our results indicate that machine learning models produce similar results to linear discriminant analysis, but in some cases machine learning produces more consistent accuracy between the sexes. Our study shows that sex can be accurately predicted (> 80%) in Italian populations using the cranial measurements we gathered, except for the foramen magnum, which shows a level of accuracy of ∼70% accurate which is on par with previous geometric morphometrics studies using crania in sex estimation. We also find that our trained machine learning models produce population-specific results; we see that Italian crania are sexually dimorphic, but the features that are important to this dimorphism differ between the populations.

## 1 INTRODUCTION

Sex estimation is an important step in identification of skeletal remains, both in bio-archaeology and forensics (Bašić et al., 2013; Cattaneo, 2007; Uytterschaut, 1986). In bio-archaeology, reconstructing the history and environment of an individual is fundamental. If the characteristics of an individual that may correlate to social variables (i.e. status), such as sex, can be identified, this provides valuable anthropological information about that individual (Novotný, 1986). Likewise, in forensics, sex estimation is a vital step in building a biological profile that may help to identify the victim of a violent crime, and help to shed light on the cause of death (Cattaneo, 2007).

In the early 1960s researchers started to investigate if the systematic measurement of bones could lead reliable sex estimation (Birkby, 1966; Novotný, 1986). One of the first used was the cranium (Giles & Elliot, 1963), probably because of its popularity in studies on human evolution, or its utility in racial (morphological race) estimation (Birkby, 1966; Crichton, 1966; Giles & Elliot, 1962). Using crania to estimate sex is useful in many forensic and anthropological contexts, such as when the remains are fragmented or displaced and the cranium may be the first (or only) remains found, due to scavenger activity (Richardson, 2004). Discriminant analysis is a solid method for sex estimation, (Giles & Elliot, 1963), but this method is often limited by sample size, and sensitive to the presence of outliers (Acuña & Rodríguez, 2005). Thus, the strive for higher accuracy in sex estimation is ongoing. Presently, some studies have peaked to over 90% accuracy in ancestry estimation, using new approaches such as geometric morphometrics, but these methods as applied to sex estimation have yielded a much lower accuracy of ∼70% (Fortes de Oliveira et al., 2012; Jiménez-Arenas & Esquivel, 2013; Murphy & Garvin, 2018; Webster & David Sheets, 2010; Zelditch, Swiderski, & David Sheets, 2012). Our study however harnesses an innovative method—machine learning (Bejdová, Dupej, Krajíček, Velemínská, & Velemínský, 2018; Bewes, Low, Morphett, Pate, & Henneberg, 2019; Gao, Geng, & Yang, 2018).

Machine learning uses a variety of algorithms that iteratively learn from data to describe that data, improve their methods, and predict outcomes. It is an application of artificial intelligence (AI) that through learning is able to perform specific tasks without being explicitly programmed (Samuel, 2000). In such methods, usually the implementer trains the model with some initial data, and after enough training data has been given, the model is able to discern the data itself. The use of machine learning is growing in the field of physical anthropology, and has proved useful (du Jardin, et al., 2009; McBride et al., 2001; Navega, et al., 2015; Santos, Guyomarc’h, & Bruzek, 2014). Indeed, a recent study used a trained model to identify the sex in Han Chinese, using CT scans of their skull, reaching an accuracy of 98% (Gao et al., 2018). Likewise, another study trained a model using over 900 CT scans, reaching over 95% accuracy (Bewes et al., 2019). Such techniques however, require expensive CT scans, where discriminant analysis requires only osteometric measurements (Martin & Saller, 1956), making the latter more prevalent in physical anthropology. These same measurements used in traditional discriminant analysis can be used to train machine learning models to determine whether these methods improve sex estimation accuracy. A recent study used x-ray osteometric measurements, from hand bones, using several different machine learning models, resulting in a method with over 80% accuracy in sex estimation for a specific age group (Darmawan, Yusuf, Kadir, & Haron, 2015). In this study, we use two Italian populations to investigate the viability of using machine learning with craniometric data to estimate sex.

## 2 METHODS

### 2.1 Populations and skeletal samples

The skeletal material used in this study was selected from the identified collections of the Anthropological Museum of the University of Bologna, Bologna, Italy. The populations we examined consist of individuals who lived in Italy between 1814 and 1922, and died between 1898 and 1944. The samples used in this study belong to two collections, the Certosa collection and the Sassari collection (Belcastro et al., 2017; Facchini, Mariotti, Bonfiglioli, & Belcastro, 2006). In the Certosa collection the cause of death and occupation are known for over 90% of the individuals, additionally over 90% of the individuals were born in the Bologna province. In the Sassari collection, the cause of death and occupation are also known for over 90% of the individuals, and over 90% of the individuals were also born in Sardinia. The sex is also known for all individuals. Individuals that had a history of trauma or dysmorphism were excluded. The crania used in both collection were all adults, and the age range was similar across sexes (18 to 70 years of age). A wide range of measurements was taken on the cranium resulting in 17 cranial measurements. Only the best-preserved materials were selected for the analysis, thus leading to 98 crania from Bologna, and 40 crania from Sardinia. Additionally, most crania have an intact foramen magnum, and therefore we were able to measure it for 212 Sardinians, and 180 Bolognese. From the same collection, we took 40 femora for each population. All samples are equally distributed among sexes. Further details about the collection are in the Supplementary Information.

### 2.2 Measurements

All craniometric measurements were taken using spreading calipers (0.01 mm). We selected 17 craniometric measurements that are common in sexual dimorphism studies (Dayal, Spocter, & Bidmos, 2008; Fortes de Oliveira et al., 2012; Jaskulska, 2005; Jiménez-Arenas & Esquivel, 2013; Uytterschaut, 1986). The measurements, including the foramen magnum, were performed according to standard protocols set out in Martin and Saller (1956). The cranial measurements are coded from M1-M63, and we chose 17 of those standard measurements to include in our study. See the Supplementary Information for more details on these measurements. To calculate the intra-observer error, the measurements of several crania were taken again by a different observer several weeks after the initial measurement. We then calculated the standard deviation for each measurement and found an average error of 0.62mm and 0.74mm in females and males respectively.

### 2.3 Linear Discriminant Analysis

Since we knew the sex of all samples, they were ideal to test discriminant functions, and compare those to our machine learning models. We used the function *lda()* of the *MASS* package on R to perform a linear discriminant analysis (LDA), and calculated the discriminant functions for all of our datasets. We split the original datasets in two subsets, using one to calculate the discriminant function, and the other to test its accuracy. The script we used can be found on github at the following link: https://github.com/earthwall/CraniaML. For more information about the discriminant functions, see the Supplementary Information.

### 2.4 Model training

In order to find the best algorithm to train our models, we used all available machine learning algorithms in Matlab 2018a. These models include decision tree learning, support-vector machines (SVM), k-nearest neighbor algorithm (KNN), and ensemble methods (Altman, 1992; Cortes & Vapnik, 1995; Fisher, 1936; Quinlan, 1986). We present the results of the models with the highest classification accuracy. To validate the models we used k-fold cross validation, which divides the dataset into k subsets of equal size, we specified two: one of them is used to train the predictor, the other is used to verify the accuracy of the predictor and vice versa. The overall accuracy of the model is computed from both subsets (Kohavi & Others, 1995). For the MATLAB code, see the Supplementary Information.

## 3 RESULTS

### 3.1 Sexual dimorphism of Italians

Overall, the cranial sexual dimorphism in Bolognese and Sardinians is similar (**Fig.1*A***). In fact, regardless of the population, most measurements are shorter in females compared to males. Some measurements however, such as cranial length (M1) and basibregmatic height (M17), are dimorphic in females from Bologna only. Although, the distribution of the foramen magnum measurements shows a high level of variability, the sexual dimorphism of these samples is consistent with measurements from the rest of the cranium, and as such is dimorphic between men and women across both populations (**Fig.1*B***).

**Figure 1.**
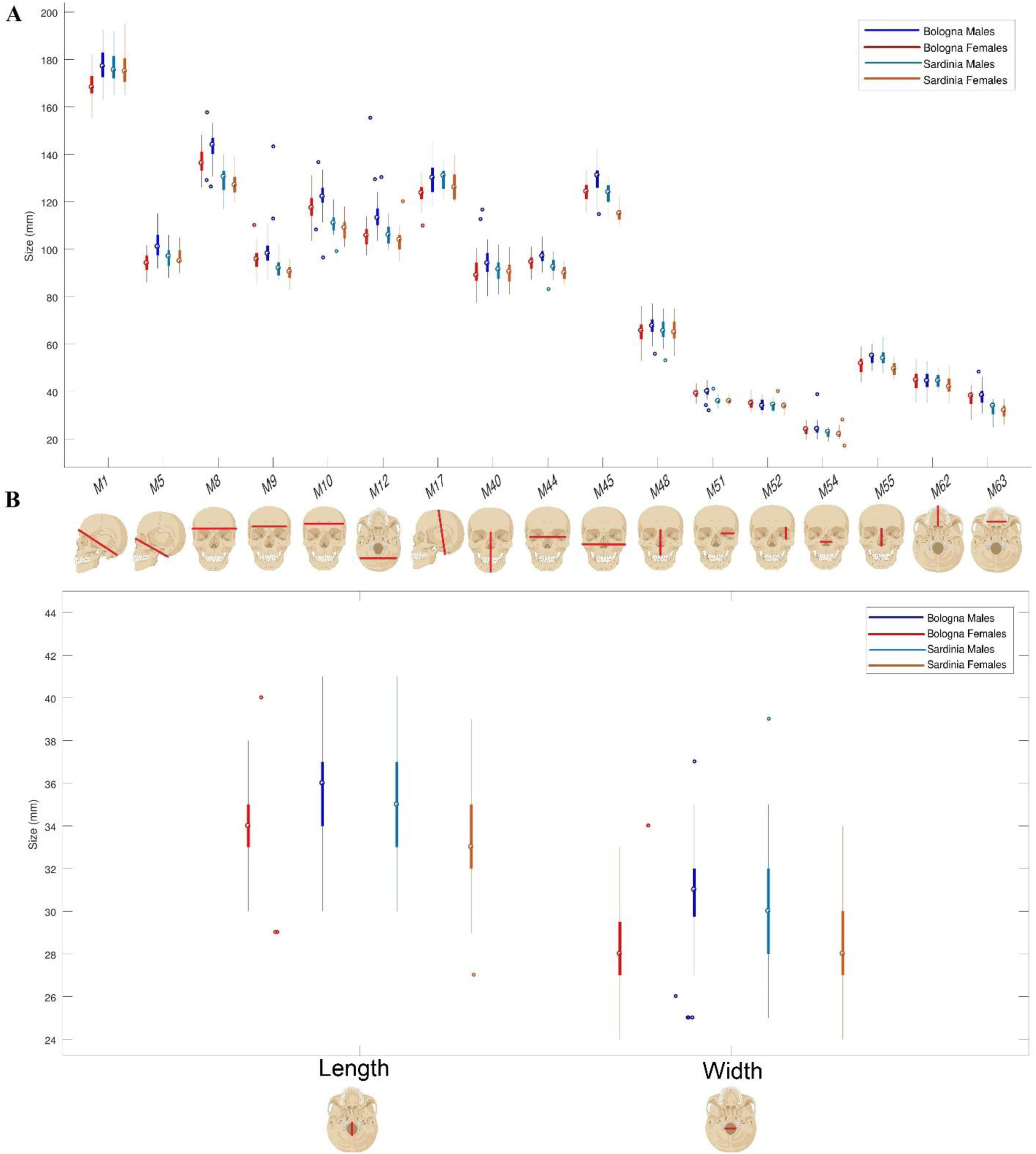
A representation of the sexual dimorphism across Italian continentals (from Bologna) and Italians from Sardinia. **A)** The boxplots represent the distribution of values for 17 cranial measurements. A representation of the section measured is present below each measurement. **B)** The boxplots represent the distribution of values for the length and width of the foramen magnum. A representation of the section measured is present below each measurement.

### 3.2 Discriminant features in Sardinians and Bolognese

Our algorithm shows us which cranial measurements are important in predicting sexually dimorphic features. The results (**Fig.2*A***) show, that based on our algorithm Sardinians and Bolognese have different features that predict sexual dimorphism. While Sardinian cranial dimorphism relies mostly on only two measurements, the sexual dimorphism in Bolognese is drawn from a wider variety of measurements. Sardinian dimorphic features are extracted mostly from the facial measurements of bizygomatic facial breadth (M45), and nasal height (M55). Instead, a variety of features are important in determining the sexual dimorphism of Bolognese, the most important of which are all linked to cranium size, such as basicranial length (M5), cranial breadth (M8), and biasteric breadth (M12).

**Figure 2.**
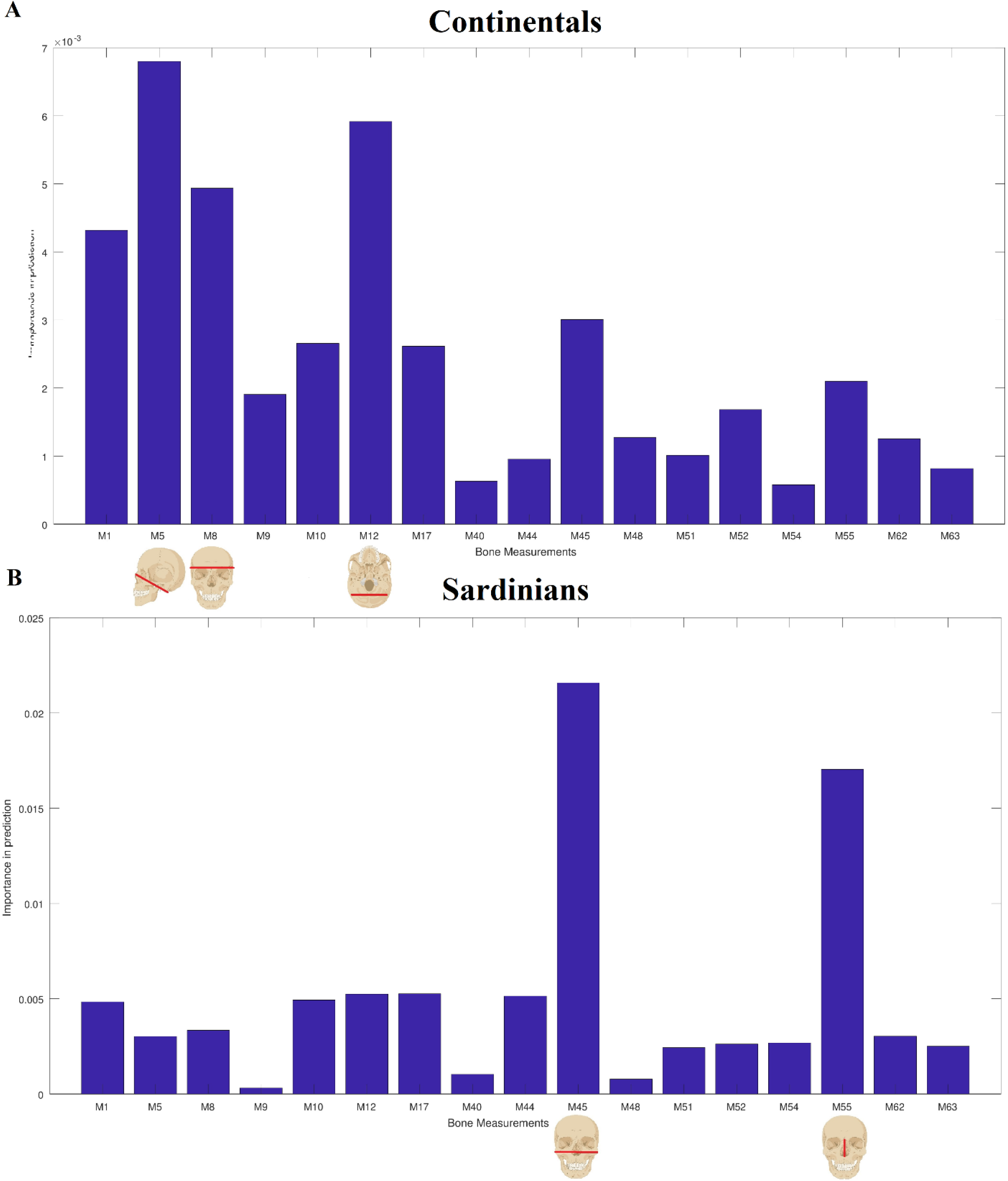
The relative importance of each measurement in the identification of sex for Italian continentals (from Bologna) and for Sardinians. The importance is shown for Italian Continentals in **(A)** and Sardinians in **(B)**.

Our results show that cranium size differs more across sexes in Bolognese than it does in Sardinians (**Fig.2*B***). We can observe that all measurements linked to cranium size (M1 to M17) are very dimorphic in Bolognese (*p* < 0.005). Conversely, in Sardinians, statistical tests (Mann-Whitney U) show an absence of any meaningful difference (*p* > 0.5) in measurements related to cranium size. However, the two populations show similarity among the other cranium measurements that are not related to cranial size, with the exception of facial breadth (M45) which shows stronger dimorphism in Sardinians. Descriptive statistics of the measurements, along with the origin of the skeletal remains, can be found in Supplementary Information.

### 3.3 Machine learning and linear discriminant analysis - Cranium

We compared the machine learning models with linear discriminant analysis for all cranial measurements except the foramen magnum. Our results show that both populations have similar results across all methods (**Tab.1**). The LDA achieved similar accuracy to the machine learning methods in the Bologna dataset, whereas accuracy was very low in the Sardinia dataset using LDA. All machine learning algorithms were better at classifying males. Furthermore, using machine learning, the misclassification of sex seems to occur mostly when the variability of a particular measurement is higher (**Fig.3**). In fact, the Sardinian sample shows less variability across sexes, and only individuals with borderline values are incorrectly classified. Conversely, the Bolognese show a higher variability of their discriminant measurements, thus increasing the amount of errors.

**Table 1.** The accuracy of the best performing machine learning methods and the linear discriminant analysis using 17 cranial measurements for both Bologna and Sardinia populations.

| Method | Population | Sex | N | Accuracy | Overall Accuracy |
| --- | --- | --- | --- | --- | --- |
| Model training: Decision tree learning | Bologna | M | 49 | 82% | 81% |
|  |  | F | 49 | 79.5% |  |
| Model training: Decision tree learning | Sardinia | M | 20 | 78% | 75% |
|  |  | F | 20 | 73% |  |
| Model training: K Nearest Neighbor | Bologna | M | 49 | 84% | 84.5% |
|  |  | F | 49 | 85% |  |
| Model training: K Nearest Neighbor | Sardinia | M | 20 | 89% | 85.5% |
|  |  | F | 20 | 82% |  |
| Linear Discriminant Analysis | Bologna | M | 49 | 83.3% | 83.3% |
|  |  | F | 49 | 83.3% |  |
| Linear Discriminant Analysis | Sardinia | M | 20 | 20% | 40% |
|  |  | F | 20 | 60% |  |

**Figure 3.**
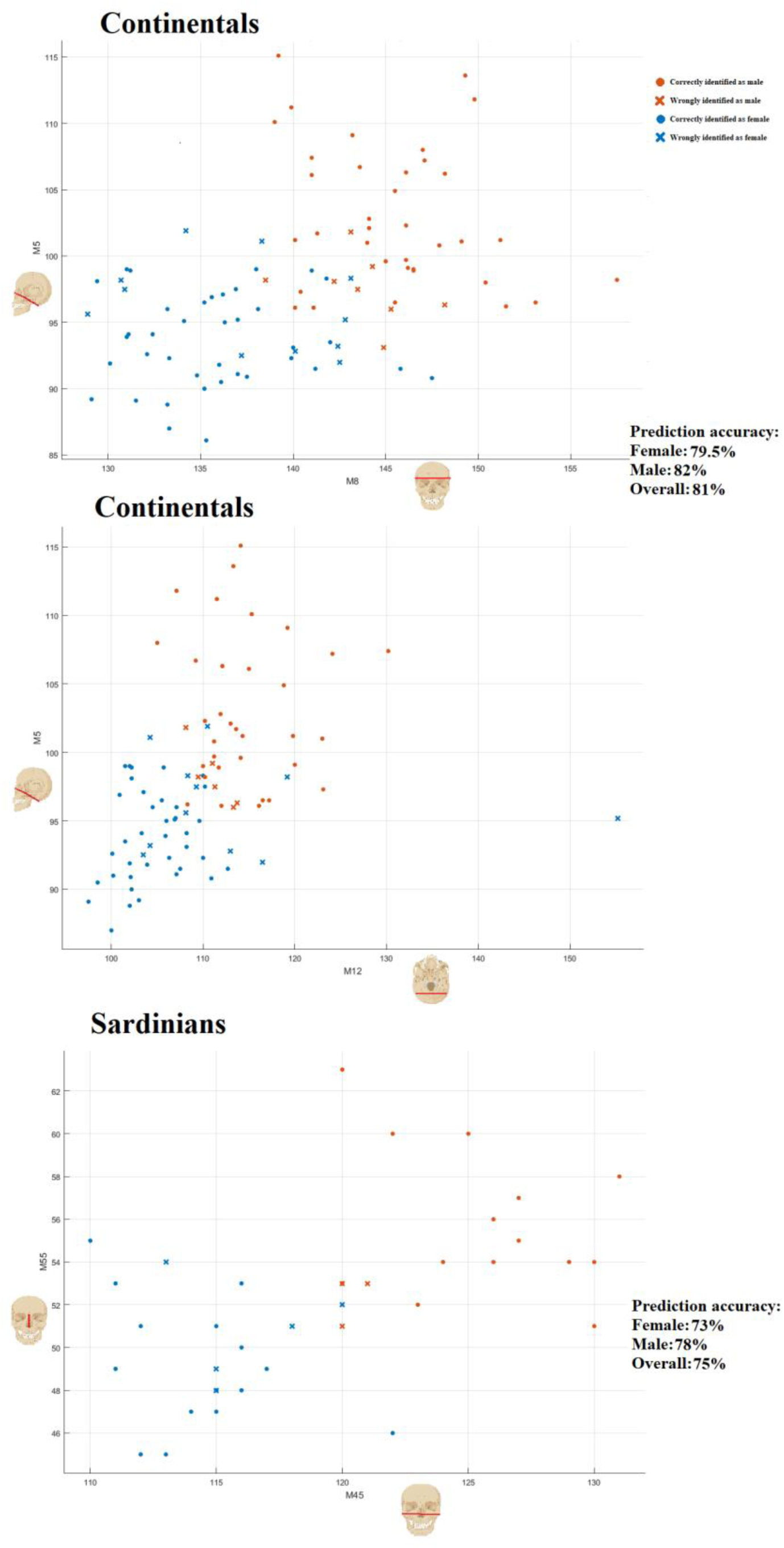
A representation of the most important measurements, along with the accuracy of the trained machine learning models. In each plot, each datum represents one individual. The individual’s identification and sex is represented by the use of different markers and colour. The colours used are orange and blue, representing males and females respectively.

The models trained using the KNN method outperformed the LDA in both populations, although only by ∼1-2%. However, KNN methods did not increase the accuracy of sex estimation equally in both sexes. While Bolognese have almost homogenous accuracy, the accuracy in Sardinians differs 7% between sexes. Therefore, the overall accuracy of sex estimation in Sardinians is higher than Bolognese by 1%, the model only performs in classifying Sardinian males (accuracy of 89%). The results produced using decision tree learning were comparable to that of the LDA methods in determining sex in the Bolognese, but the LDA still outperformed the decision tree algorithm (accuracy of 81% compared to 83.3% overall). The decision tree algorithm was much less accurate at classification of sex in the Sardinian sample (accuracy of 75% overall). Furthermore, even though the decision tree algorithm produced an accuracy of 81% overall for Bolognese, there is a large difference in accuracy between the sexes (82% for males and 79.5% for females). Therefore the most accurate method overall is the KNN algorithm.

We tested the ability of the machine learning models and the LDA to work on populations different from the ones used for the training. We used the models trained with Bolognese data to estimate sex in the Sardinian dataset and vice versa. This analysis however, gave a very low accuracy (< 50%), suggesting that these algorithms will only produce a high level of accuracy when used within the original population used to train them. Likewise, the discriminant functions calculated through the LDA had very skewed accuracy with overall very low accuracy (< 50%). This suggests that the variability in cranial measurements is too high to be reliably used in sex prediction even across populations that are relatively close, such as Bologna and Sardinia. Likewise, the circles and crosses, represent correctly and incorrectly identified individuals respectively.

### 3.4 Machine learning and linear discriminant analysis - Foramen Magnum

Both the machine learning methods and the LDA performed similarly for both populations when applied to the foramen magnum dataset (accuracy of ∼70%) (**Tab.2**). The accuracy was similar regardless of sample size, indeed when using only 50% of the Sardinian dataset to compute a discriminant function the accuracy of the prediction dropped by only 1%. Nonetheless, the accuracy of the sex estimation is quite different between the sexes, across both methods, and across all the datasets. Using machine learning methods, the accuracy varies by 10% between Sardinian males and females, and 6% for Bolognese. Using the linear discriminant analysis the functions computed have a difference in accuracy between the sexes of ∼7% in the Bologna dataset, ∼6% in the Sardinia dataset, and over 17% in the mixed dataset. We reported here only the results of the best performing models, however most models had unbalanced accuracy between the sexes (data not shown). As these differences are consistent across methods, the discrepancies in accuracy for each sex are probably not related to using different learning methods for each population. Rather, males have usually longer and wider foramen magnums compared to females in both populations, and probably the methods are sensitive to these differences (**Fig.4**). Indeed, most misclassified males are the ones with a smaller foramen magnum, and most misclassified females have a bigger foramen magnum. Following this rationale, both extremes, males with a large foramen magnum and females with a small one, have very few misclassifications, this pattern is consistent across both populations. Nonetheless, machine learning methods outperformed traditional methods in the Sardinian sample.

**Table 2.** The accuracy of the best performing machine learning methods and the linear discriminant analysis using foramen magnum measurements for both Bologna and Sardinia populations.

| Method | Population | Sex | N | Accuracy | Overall Accuracy |
| --- | --- | --- | --- | --- | --- |
| Model training: Support Vector Machine | Bologna | M | 80 | 69% | 72% |
|  |  | F | 80 | 75% |  |
| Model training: Support Vector Machine | Sardinia | M | 106 | 75% | 70% |
|  |  | F | 106 | 65% |  |
| Model training: Quadratic Discriminant | Mixed | M | 186 | 68% | 70.2% |
|  |  | F | 186 | 72% |  |
| Linear Discriminant Analysis | Bologna | M | 80 | 65% | 73.7% |
|  |  | F | 80 | 82% |  |
| Linear Discriminant Analysis | Sardinia | M | 106 | 58% | 61.3% |
|  |  | F | 106 | 64% |  |
| Linear Discriminant Analysis | Mixed | M | 80 | 80% | 71.4% |
|  |  | F | 80 | 62.5% |  |

**Figure 4.**
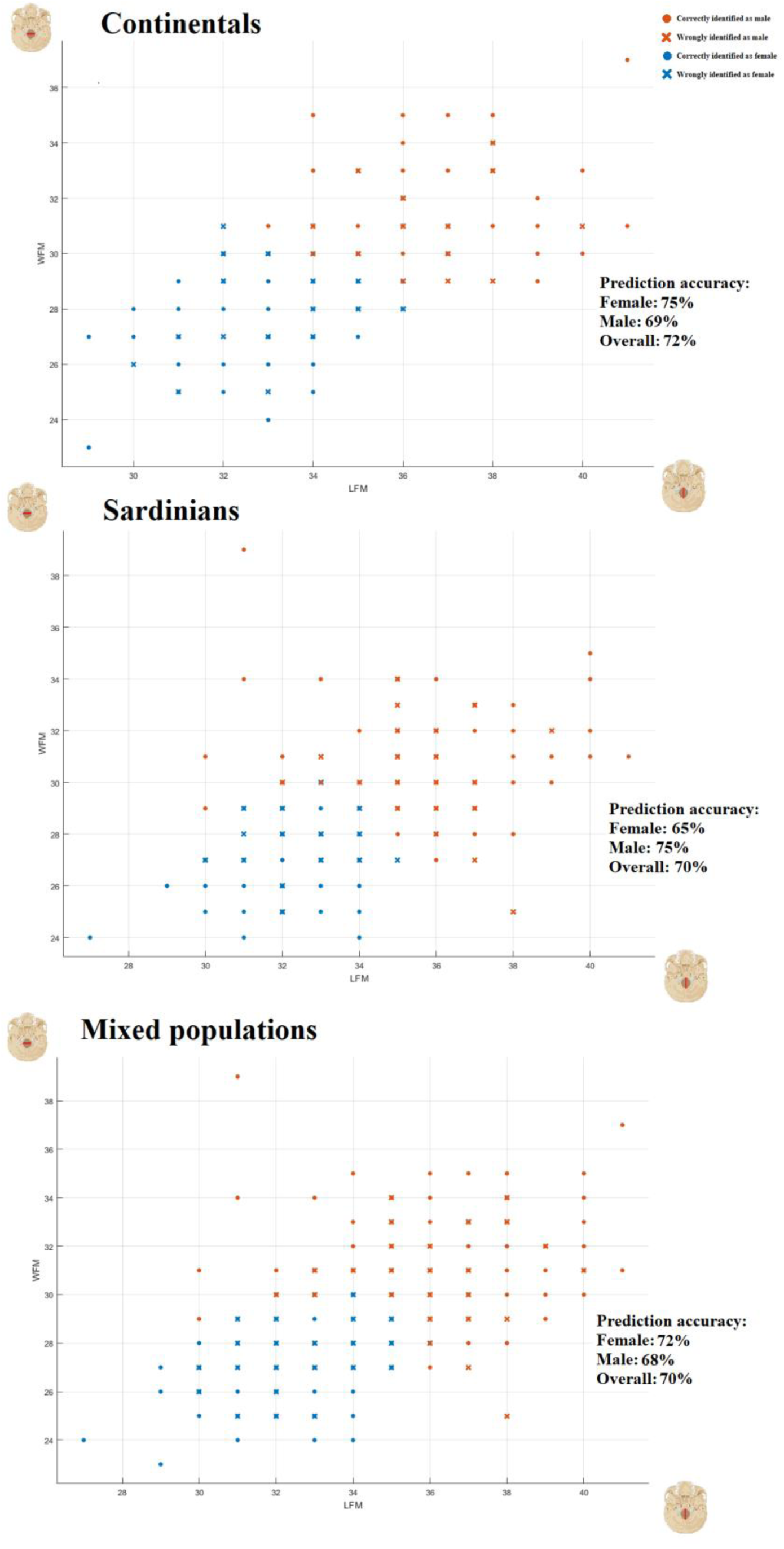
Representation on the axis of length and width of the foramen magnum, along with the accuracy of the trained machine learning models. In each plot, each datum represents one individual. The individual’s identification and sex is represented by the use of different markers and colour. The colours used are orange and blue, representing males and females respectively. Likewise, the circles and crosses, represent correctly and incorrectly identified individuals respectively.

## 4 Discussion

Here we present results for sex estimation using cranial measurements from two Italian populations. We compare the use of machine learning methods and traditional methods and find that while machine learning and linear discriminant analysis often have similar accuracy, the machine learning models show accuracy that is more consistent. Nonetheless, both of these methods failed in creating reliable algorithms, or functions, in predicting sex in a new dataset. In fact, regardless of whether we used machine learning or linear discriminant analysis, each method was accurate only with individuals from the same population. Interestingly, we provide further evidence that the foramen magnum is useful for the estimation of sex (Raikar, Meundi, David, Rao, & Jogigowda, 2016). Indeed, the methods applied to the foramen magnum measurements yielded a consistent 70% accuracy across populations. The trained models have similar accuracy on both sexes when applied to multiple populations, compared to almost 20% difference in the linear discriminant method, suggesting that the machine learning methods might be more suitable when using these measurements. We add to an emerging and yet growing body of knowledge that suggests machine learning (in combination with craniometry) is useful in estimating sex, albeit results are population specific, also a limitation of more traditional methods (du Jardin, et al., 2009; McBride et al., 2001; Murphy & Garvin, 2018; Navega, et al., 2015; Santos, Guyomarc’h, & Bruzek, 2014).

## Acknowledgments

We thank Matteo Venturini and Serena Cappa for their help in taking the measurements necessary for the calculation of the intra-observer error.

## Data Availability Statement

Data and codes used for this paper are available in the supplementary information and on github https://github.com/earthwall/CraniaML; if you require further information please contact the corresponding author

## Author Contributions

AP and CR designed the study. AP and CR took the measurements of the samples. AP performed all the analyses. AP, TLC, CR, and MB interpreted the results. TLC and AP wrote the manuscript.

## Supplementary Information

The supplementary information includes four tables and one figure and provides more information on the study.

The four tables have several pieces of information on the measurements taken:

Table S1: Bologna cranial measurements (including mean and number of samples per sex)

Table S2: Sardinian cranial measurements (including mean and number of samples per sex)

Table S3: Foramen magnum measurements (including mean and number of samples per sex)

Table S4: Origin of the crania used for the foramen magnum measurements.

Figure S1: The figure represents the path of choices made by an AI trained with decision tree learning method.

Lastly, we report the discriminant function computed using the *MASS* package of R in the last page of the SI.

**Table S1:** Bologna cranial measurements. In the following table several information for 17 different osteocranial measurements are reported. The information includes the common name for the variable, the reference number according to Martin & Saller, the name of the landmarks used for the measurement, sex, the number of individuals in which we were able to take the measurement, the average, and the standard deviation. All measurements are in millimetres. In the last column we report the p-value on a one tailed t-student test between males and females. The result of the test is approximated to the 4^th^ decimal.

| **Variable** | **Martin no.** | **Measurement Landmarks** | **Sex** | **N** | **mean** | **s.d.** | **p** |
| --- | --- | --- | --- | --- | --- | --- | --- |
| Cranial length | M1 | Glabella -Opisthocranion | M<br>F | 49<br>46 | 177.33<br>168.62 | 6.46<br>5.97 | 0 |
| Basicranial length | M5 | Basion-Nasion | M<br>F | 48<br>48 | 101.43<br>94.183 | 5.61<br>3.58 | 0 |
| Cranial breadth | M8 | Eurion - Eurion | M<br>F | 49<br>48 | 143.40<br>136.94 | 6.17<br>5.25 | 0 |
| Minimum frontal breadth | M9 | Frontotemporale<br>Frontotemporale | M<br>F | 49<br>47 | 99.36<br>96.17 | 7.88<br>4.28 | 0.0065 |
| Maximum frontal breadth | M10 | Coronale - Coronale | M<br>F | 48<br>48 | 121.82<br>117.47 | 6.80<br>5.94 | 0.0006 |
| Biasteric breadth | M12 | Eurion - Eurion | M<br>F | 46<br>45 | 114.73<br>105.46 | 8.45<br>4.15 | 0 |
| Basibregmatic height | M17 | Basion - Bregma | M<br>F | 47<br>48 | 129.95<br>123.83 | 7.07<br>4.60 | 0.0001 |
| Facial length | M40 | Basion - Prosthion | M<br>F | 48<br>48 | 94.39<br>89.98 | 6.95<br>5.21 | 0.0004 |
| Biorbital breadth | M44 | Ectonochion - Ectonochion | M<br>F | 47<br>44 | 97.15<br>94.14 | 3.35<br>3.67 | 0 |
| Bizygomatic breadth | M45 | Zygion-Zygion | M<br>F | 44<br>38 | 129.77<br>124.11 | 5.52<br>4.05 | 0 |
| Superior facial height | M48 | Nasion-Prosthion | M<br>F | 49<br>48 | 67.99<br>65.45 | 4.71<br>5.06 | 0.0087 |
| Orbital breadth (R) | M51 | Dacryon-Ectonochion | M<br>F | 48<br>46 | 39.92<br>39.23 | 2.43<br>1.94 | 0.0553 |
| Orbital height (R) | M52 | No Landmarks | M<br>F | 48<br>46 | 34.35<br>35.17 | 2.33<br>2.42 | 0.0536 |
| Nasal breadth | M54 | Alare - Alare | M<br>F | 48<br>46 | 24.43<br>23.84 | 2.64<br>2.05 | 0.1172 |
| Nasal height | M55 | Nasion-Nasospinale | M<br>F | 49<br>47 | 54.46<br>51.43 | 2.88<br>3.81 | 0 |
| Palate length | M62 | Prosthion – Alveolar border | M<br>F | 48<br>49 | 44.48<br>44.80 | 4.03<br>4.19 | 0.3455 |
| Palate breadth | M63 | Endomolare – Endomolare | M<br>F | 48<br>49 | 38.17<br>37.09 | 3.47<br>3.61 | 0.0694 |

**Table S2:** Sardinia cranial measurements. In the following table several information for 17 different osteocranial measurements are reported. The information includes the common name for the variable, the reference number according to Martin & Saller, the name of the landmarks used for the measurement, sex, the number of individuals in which we were able to take the measurement, the average, and the standard deviation. All measurements are in millimetres. In the last column we report the p-value on a one tailed t-student test between males and females. The result of the test is approximated to the 4^th^ decimal.

| Variable | Martin no. | Measurement Landmarks | Sex | N | mean | s.d. | p |
| --- | --- | --- | --- | --- | --- | --- | --- |
| Cranial length | M1 | Glabella -Opisthocranion | M<br>F | 20<br>20 | 176.65<br>175.3 | 6.82<br>7.24 | 0.2788 |
| Basicranial length | M5 | Basion-Nasion | M<br>F | 20<br>20 | 96.5<br>96.2 | 4.2<br>3.92 | 0.4106 |
| Cranial breadth | M8 | Eurion - Eurion | M<br>F | 20<br>20 | 129.25<br>127.6 | 5.19<br>4.5 | 0.151 |
| Minimum frontal breadth | M9 | Frontotemporale<br>Frontotemporale | M<br>F | 20<br>20 | 92.4<br>90.4 | 3.84<br>3.34 | 0.0474 |
| Maximum frontal breadth | M10 | Coronale - Coronale | M<br>F | 20<br>20 | 110.95<br>108.5 | 4.55<br>4.78 | 0.057 |
| Biasteric breadth | M12 | Eurion - Eurion | M<br>F | 20<br>20 | 106.3<br>104.2 | 4.18<br>5.21 | 0.0894 |
| Basibregmatic height | M17 | Basion - Bregma | M<br>F | 20<br>20 | 129.85<br>127.4 | 4.77<br>6.32 | 0.0927 |
| Facial length | M40 | Basion - Prosthion | M<br>F | 20<br>20 | 91.5<br>90.6 | 5.63<br>5.2 | 0.3059 |
| Biorbital breadth | M44 | Ectonochion - Ectonochion | M<br>F | 20<br>20 | 92.7<br>89.9 | 3.9<br>2.9 | 0.0082 |
| Bizygomatic breadth | M45 | Zygion-Zygion | M<br>F | 20<br>20 | 123.25<br>115.3 | 5.17<br>3.32 | 0 |
| Superior facial height | M48 | Nasion-Prosthion | M<br>F | 20<br>20 | 66.2<br>65.25 | 5.31<br>4.93 | 0.2854 |
| Orbital breadth (R) | M51 | Dacryon-Ectonochion | M<br>F | 20<br>20 | 36.3<br>36.05 | 1.98<br>1.36 | 0.3262 |
| Orbital height (R) | M52 | No Landmarks | M<br>F | 20<br>20 | 34.2<br>33.95 | 2.06<br>2.18 | 0.3593 |
| Nasal breadth | M54 | Alare - Alare | M<br>F | 20<br>20 | 22.3<br>22.3 | 1.73<br>2.35 | 0.5 |
| Nasal height | M55 | Nasion-Nasospinale | M<br>F | 20<br>20 | 54.45<br>49.6 | 3.65<br>2.87 | 0 |
| Palate length | M62 | Prosthion – Alveolar border | M<br>F | 20<br>20 | 44.65<br>42.8 | 3.13<br>4.34 | 0.0702 |
| Palate breadth | M63 | Endomolare – Endomolare | M<br>F | 20<br>20 | 33.2<br>31.55 | 3.28<br>3.11 | 0.0598 |

**Table S3:** Foramen Magnum measurements. In the following table several information are reported on the foramen magnum measurements. The measurements are in millimetres. In the last column we report the p-distance on a one tailed tstudent test between males and females. The result of the test is approximated to the 4^th^ decimal.

| Variable | Population | Sex | N | mean | s.d. | p |
| --- | --- | --- | --- | --- | --- | --- |
| Foramen Magnum Length | Bologna | M | 80 | 35.81 | 2.49 | 0 |
|  |  | F | 80 | 33.87 | 2.24 |  |
| Foramen Magnum Width | Bologna | M | 80 | 30.65 | 2.33 | 0 |
|  |  | F | 80 | 28.45 | 2.10 |  |
| Foramen Magnum Length | Sardinia | M | 106 | 35.05 | 2.52 | 0 |
|  |  | F | 106 | 33.39 | 2.05 |  |
| Foramen Magnum Width | Sardinia | M | 106 | 30.11 | 2.24 | 0 |
|  |  | F | 106 | 28.22 | 2.10 |  |

**Table S4:**
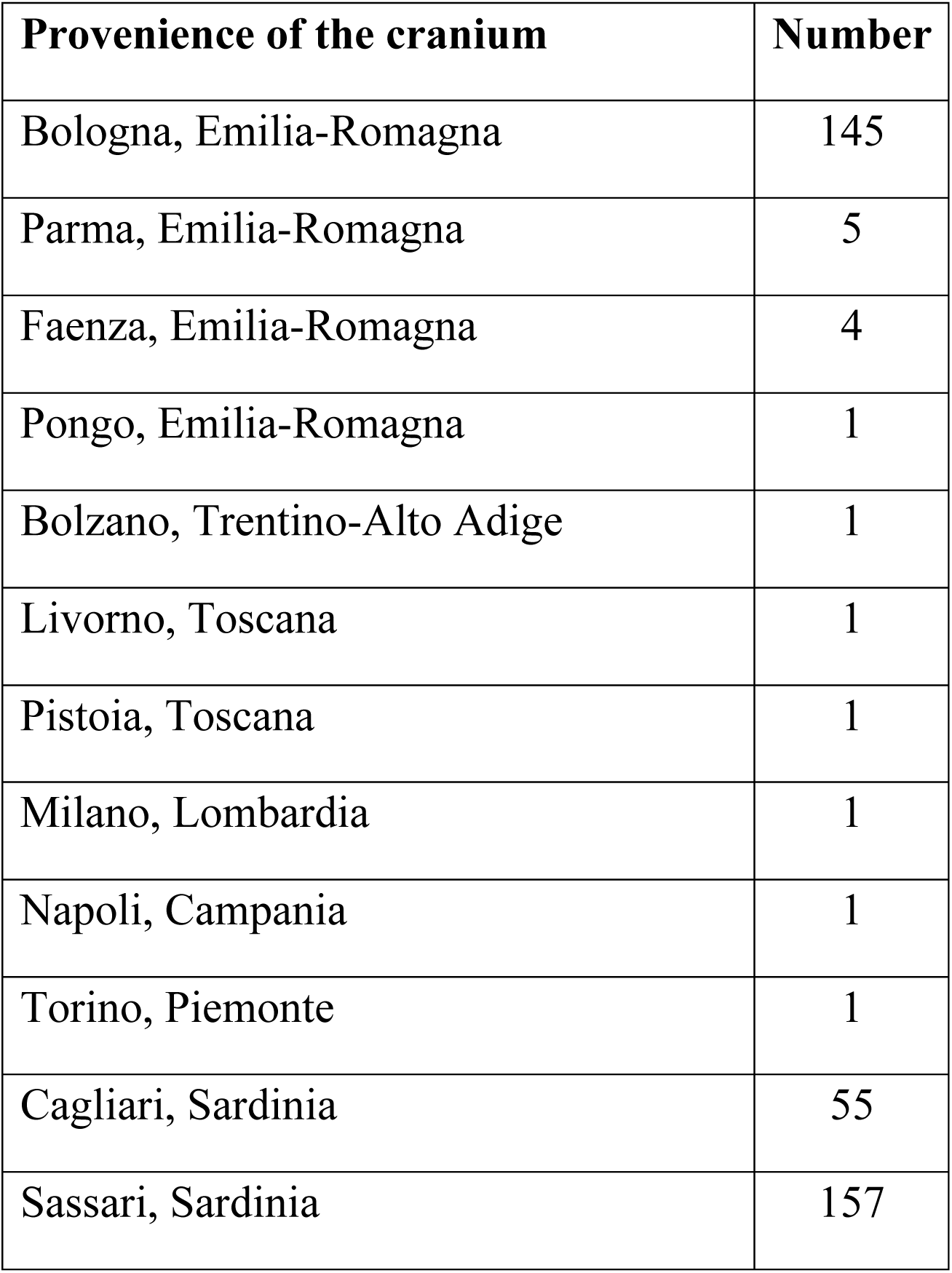
Foramen Magnum – Cranium origin. In the following table, we listed the provenience of the cranium used for the foramen magnum measurements. The name of the city and region of provenience are listed.

| Provenience of the cranium | Number |
| --- | --- |
| Bologna, Emilia-Romagna | 145 |
| Parma, Emilia-Romagna | 5 |
| Faenza, Emilia-Romagna | 4 |
| Pongo, Emilia-Romagna | 1 |
| Bolzano, Trentino-Alto Adige | 1 |
| Livorno, Toscana | 1 |
| Pistoia, Toscana | 1 |
| Milano, Lombardia | 1 |
| Napoli, Campania | 1 |
| Torino, Piemonte | 1 |
| Cagliari, Sardinia | 55 |
| Sassari, Sardinia | 157 |

**Figure S1:**
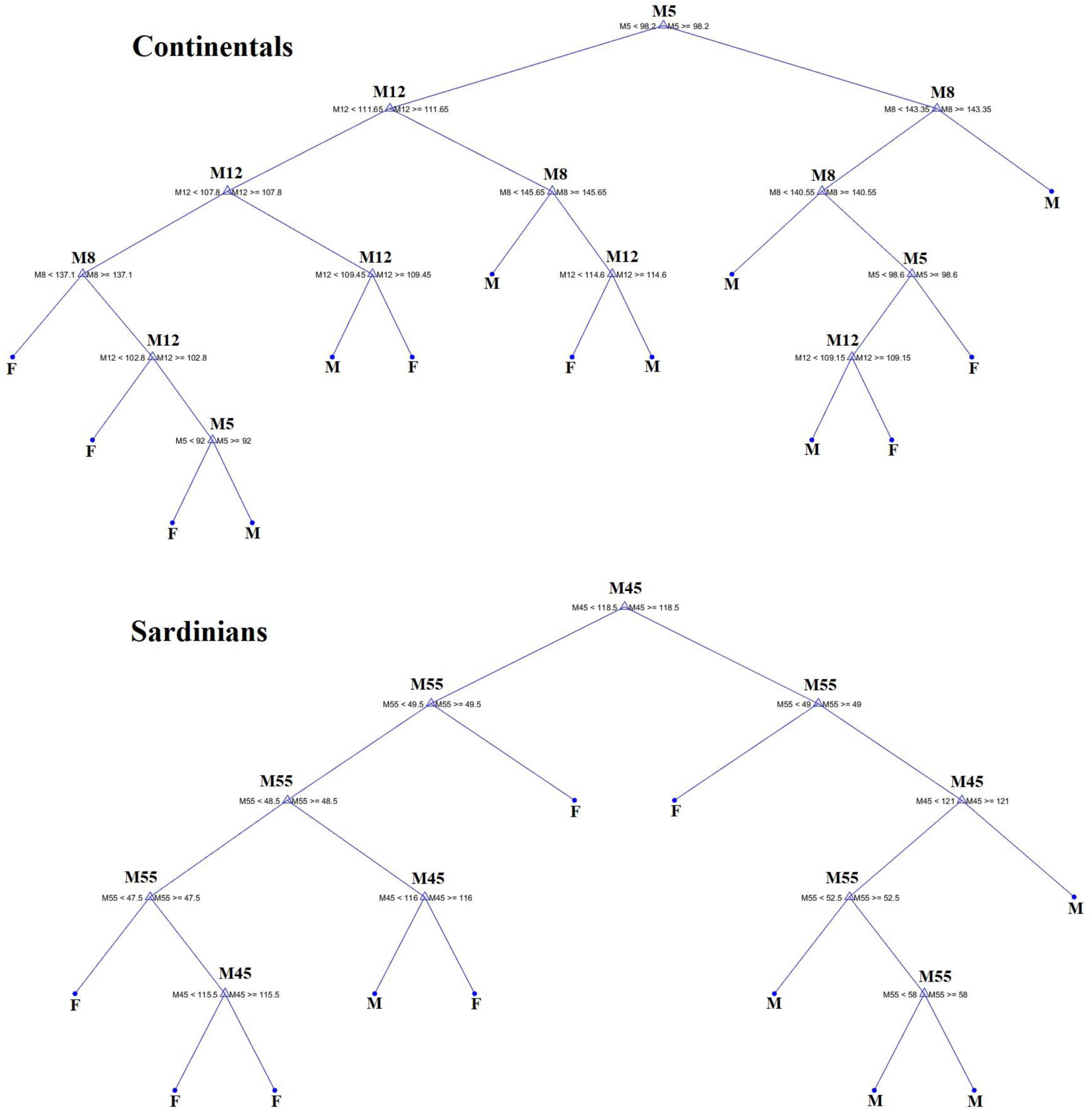
AI trained with decision tree learning method In the following figure, there is a representation of the decision tree learning method for both populations, Sardinian and continental Italians (Bologna). The trees show the number of decisions taken by the AI in the two populations, and thus shows the process used to identify the two sexes. According to the algorithm, in continental Italians the basicranial length (M5) measurement divide the algorithm into two different branches. The left branch relies mostly on measurements of the biasteric breadth (M12), however both branches use all the measurements. The algorithm in the AI classifying Sardinians splits in two branches as well, only this time using the value of bizygomatic breadth (M45). The branches divide almost perfectly into males on the left branch, and females on the right branch.

## Linear Discriminant Analysis

Here are listed the linear discriminant functions calculated in this study. We refer to individual measurements within the functions by using their Martin Code (e.g. M1), except for the foramen magnum.

This is the discriminant function for the Bologna Dataset calculated using all the craniometric measurements available. Closer the X is to 1, more is likely that the individual is female. The accuracy is 81%.

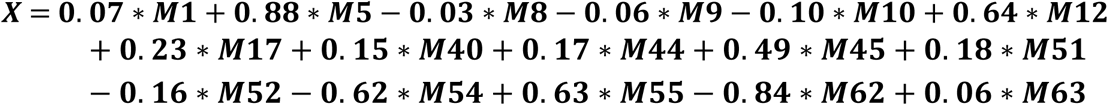

This is the discriminant function for the Bologna Dataset calculated using only the most discriminant measurements. Closer the X is to 1, more is likely that the individual is female. The accuracy is: 83%

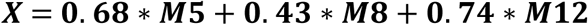

This is the discriminant function for the Sardinia Dataset calculated using all the craniometric measurements available. Closer the X is to 1, more is likely that the individual is female. The accuracy is 40%.

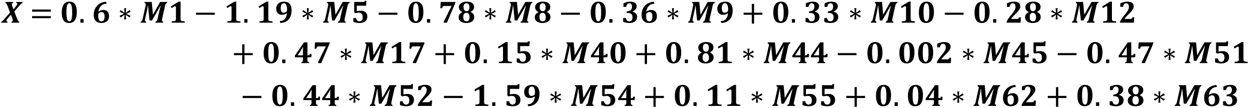

This is the discriminant function for the Sardinia Dataset calculated using only the most discriminant measurements. Closer the X is to 1, more is likely that the individual is female. The accuracy is 45%.

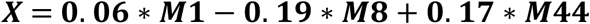

This is the discriminant function for the Bologna FM Dataset calculated using width and length of the FM. Closer the X is to 1, more is likely that the individual is female. The accuracy is 73.7%.

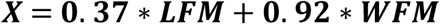

This is the discriminant function for the Sardinia FM Dataset calculated using width and length of the FM. Closer the X is to 1, more is likely that the individual is female. The accuracy is 61.3%.

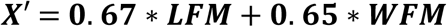

This is the discriminant functions using half (X, 106 individuals) and whole (Y, 212 individuals) of the Sardinia FM Dataset and tested using only the Bologna Dataset. This is to compare the sample size effect on the discriminant function coefficients and accuracy.

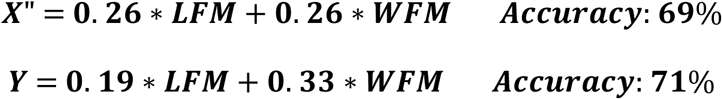

The coefficient in the two functions (X’ and X”) used for the Sardinia FM dataset are different because the partitions used to compute the function, and to test it, are random each time. However, in each case both LFM and WFM have very similar coefficients, demonstrating that the resulting function is quite consistent.

